# Whole genome analysis identifies the association of *TP53* genomic deletions with lower survival in Stage III colorectal cancer

**DOI:** 10.1101/784645

**Authors:** Li C Xia, Paul Van Hummelen, Matthew Kubit, Hojoon Lee, John M Bell, Susan M. Grimes, Christina Wood-Bouwens, Stephanie U. Greer, Tyler Barker, Derrick S Haslem, James Ford, Gail Fulde, Hanlee P Ji, Lincoln D Nadauld

**Author notes:** Corresponding authors: Hanlee P. Ji, Lincoln D. Nadauld.

## Abstract

DNA copy number aberrations **(CNA)** were frequently observed in colorectal cancers **(CRC)**. There is an urgent call for CNA-based biomarkers in clinics, in particular for Stage III CRC, if combined with imaging or pathologic evidence, promise more precise care at the timing. We conducted this Stage III specific biomarker discovery with a cohort of 134 CRCs, and with a newly developed high-efficiency CNA profiling protocol. Specifically, we developed the profiling protocol for tumor-normal matched tissue samples based on low-coverage clinical whole-genome sequencing (**WGS**). We demonstrated the protocol’s accuracy and robustness by a systematic benchmark with microarray, high-coverage whole-exome and -genome approaches, where the low-coverage WGS-derived CNA segments were highly accordant (PCC>0.95) with those derived from microarray, and they were substantially less variable if compared to exome-derived segments. A lasso-based model and multivariate cox regression analysis identified a chromosome 17p loss, containing the *TP53* tumor suppressor gene, that was significantly associated with reduced survival (P=0.0139, HR=1.688, 95% CI = [1.112-2.562]), which was validated by an independent cohort of 187 Stage III CRCs. In summary, the new low-coverage WGS protocol has high sensitivity, high resolution and low cost and the identified 17p-loss is an effective poor prognosis marker for Stage III patients.

## INTRODUCTION

Extensive copy number aberrations **(CNA)** are a hallmark of cancers with genome instability and are observed among a wide variety of epithelial malignancies originating from the colon, breast, cervix, prostate, bladder and stomach ^1^. High levels of CNAs are associated with cancer progression and poor prognosis. Thus, there is general interest in profiling CNAs as potential biomarkers associated with specific clinical outcome.

The focus of our study was colorectal cancer **(CRC)**, the third most common cancer world-wide, with ∼1.8 million estimated new cases yearly. The majority of CRCs demonstrates an extensive CNAs and as a result, are designated as belonging to the chromosomal-instability **(CIN)** molecular subtype. To accurately profile CNAs and evaluate their prognostic significance in CRC, a variety of methods have been used which include karyotyping, fluorescent in-situ hybridization **(FISH)**, and chromosomal microarrays, such as comparative genomic hybridization **(CGH)** arrays and single nucleotide polymorphisms **(SNP)** arrays.

Citing some examples, with karyotyping, *Bardi et al*. found that loss of chromosome 18 was correlated with shorter overall survival in early-stage patients (N=150) ^2^. *Personeni et al*. ^3^using FISH identified that changes in *EGFR* copy number predicted overall survival for EGFR-targeted therapy in a metastatic CRC cohort (N=87). Using SNP-arrays, *Sheffer et al*. identified deletions of 8p, 4p, and 15q were associated with poor survival in a mixed-stage cohort (N=130) ^4^. Other microarray-based studies ^5-7^ using smaller numbers of patients (N<100) identified various CNAs associated with poor survival that included sub-arm losses of 1p, 4p, 5, 6, 8p, 10, 14q, and 18. In contrast, based on CGH-array analysis, *Rooney et al*. ^8^ reported that no specific CNA was significantly associated with survival in Duke’s C-stage CRCs (N=29). The lack of concordance among these studies reflects the clinical stage variation among the study cohorts and the inherent limits of the molecular methods used for detecting CNAs and suggests clinical stage-specific variation among the study cohorts.

More recently, researchers have employed whole genome **(WGS)**, whole exome **(WES)**, and targeted sequencing for high-resolution analysis to profile CNAs. WGS has significant advantages over the other approaches because it provides whole-genome coverage without targeting and capturing as compared to other methods including microarrays and exomes. Exome and targeted sequencing have technical biases due to the extra DNA amplification and hybridization steps. Furthermore, these methods with their emphasis on gene targets cover only a small proportion (<3%) of the genome and thus miss significant portions of the noncoding genome which are noncoding. However, conducting high-coverage WGS is costly even when considering recent cost reductions in sequencing. It also generates large data sets that incur significant informatics cost.

To overcome some of the challenges of conducting cancer WGS studies on populations, we developed a low-coverage whole genome approach that provided highly accurate genome-wide copy number results. As a result, this leads to a significantly lower per sample cost than conventional WGS. For this study, we used an average sequencing coverage of less than 5X. Moreover, this WGS approach was optimized for sequencing formal fixed paraffin embedded **(FFPE)** samples, thus enabling our approach to be widely used for archival pathology biopsies. We verified the accuracy of CNA segments generated with this approach using a systematic benchmark with microarray, WES and high-coverage WGS. Overall, development of this low-coverage WGS enabled us to analyze the entire genome for CNAs while reducing the sequencing cost and bioinformatic workload.

We applied this WGS approach to a Stage III CRC cohort. Among the different clinical stages of CRC, there is a particular interest in identifying CNAs that indicate poor prognosis for individuals with Stage III disease, where local lymph node involvement is present without imaging or pathologic evidence of distal metastasis. These patients routinely receive adjuvant chemotherapy and yet, there is a significant fraction that show recurrence despite receiving adjuvant treatment after complete resection of their cancers. Identifying Stage III patients at high risk for recurrence may prove useful in targeting these individuals for more effective adjuvant regimens and developing more sensitive screening protocols for detecting early metastasis.

We conducted a WGS analysis on a discovery cohort of Stage III CRCs (N=134). We determined whether any specific CNAs were associated with a poor survival within the cohort. We validated our results with an independent cohort of Stage III CRCs from the Cancer Genome Atlas **(TCGA)** project (N=187). Our findings provided additional evidence to support that specific CNAs are predictive for CRC progression, having identified a specific genomic deletion that is predictive for lower overall survival.

## RESULTS

### Copy number calling

We benchmarked genome segmentation by bioinformatics tools such as CNVkit and Bic-seq on low-coverage WGS data with a randomly selected set of 10 tumor-normal matched CRCs from TCGA **(Fig. 1)**. Their copy number analysis data were also publicly available from SNP microarrays. Using either caller, we observed that the WGS-derived genome segments were highly correlated with the microarray segments, which were considered as ground-truth **(Fig. 2)**. The genome-wide tile-based average PCC were, 0.966 and 0.963 for CNVkit and Bic-Seq, respectively, and the gene-based ones were 0.943 and 0.938. We observed no statistically significant difference between PCC metrics of CNVkit and Bic-seq (Wilcoxon’s P=0.68). We selected CNVkit because of its consistent performance with a relatively smaller median absolute deviation (*i.e*. intra-tile deviation metrics): 0.0088 vs 0.0113.

**Figure 1.**
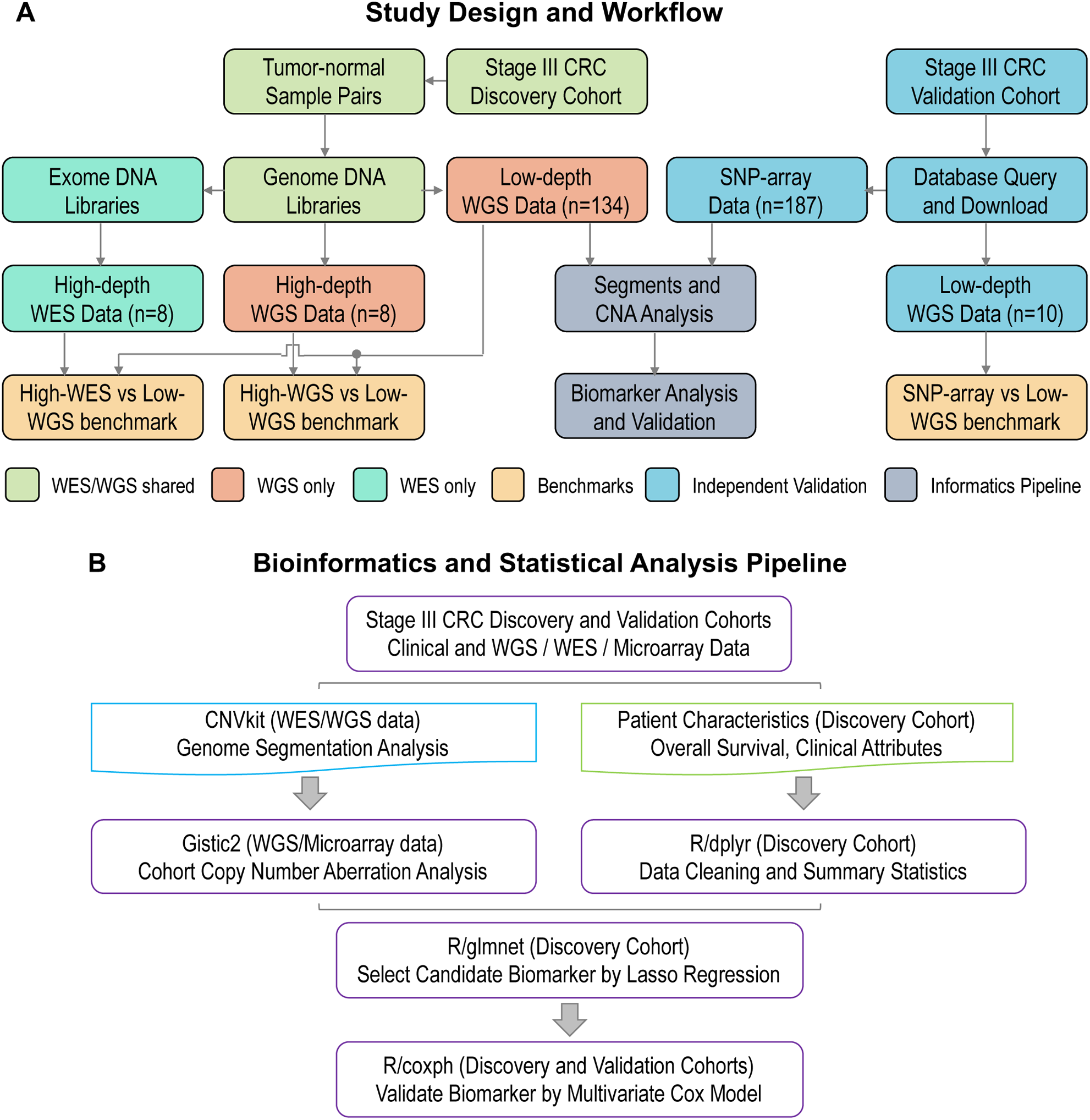
The study design and whole genome sequencing analytical workflow. **(A)** The whole-genome sequencing (WGS) analysis share the same sample preparation, DNA extraction and quality control steps as WES (color shaded light green). The prepared genomic DNA libraries are pooled for WGS directly, while they require additional PCR amplification and hybridization steps to generate exomic libraries for pooled WES. **(B)** We integrated CNVkit, Gistic2 and various R packages to perform copy number segmentation, CNA calling and biomarker discovery analyses.

**Figure 2.**
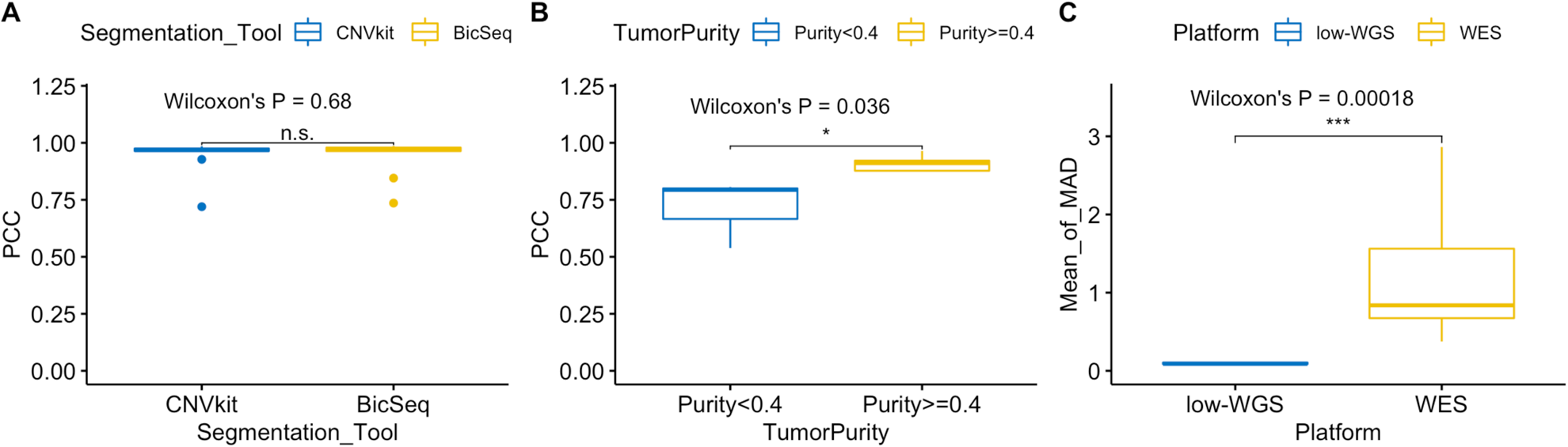
Benchmarks to evaluate low-coverage WGS approach and bioinformatics. (**A**) Pearson’s correlation coefficients (PCC) between low-coverage WGS and microarray segments as stratified by segmentation tools; (**B**) PCC between low-coverage and high-coverage WGS as stratified by tumor purity; (**C**) The means of robust standard deviation (MAD) as stratified by low-coverage WGS and WES analysis platforms.

We examined the potential variability and bias as related to low average coverage. For this evaluation, we sequenced eight pilot tumor-normal pairs at a higher coverage at ∼30x and these high-coverage WGS data were down-sampled to ∼3x. We found high correlations between low- and high-coverage WGS segments with an average PCC at 0.93 **(Fig. 2)**. We identified the only three samples that had PCC<0.9 all had tumor purity <40% (Wilcoxon’s P=0.036), suggesting that the sensitivity of detection could be limited by the poor sample quality given an adequate coverage (>2x).

We compared genome segmentation results between low-coverage WGS and WES platforms. We found a general agreement between methods although the WES CNV estimates had a higher level of noise. Within a genome tile size of 100kb, we did not generally expect abrupt copy number changes within a tile, therefore the intra-tile CNR deviation are mostly because of experimental variability as related to the sequencing preparation. We found that the mean intra-tile copy number ratio deviation (as measured by median absolute deviation) was much higher in WES derived segmentations as compared to low-coverage WGS (Wilcoxon’s P=0.00018). Specifically, the intra-tile deviation metric was as small as <0.1 for WGS while it was 1.09 for WES. Our results point to WES-derived segments being highly variable, which is apparent from visual inspection of the segmentation tracks compared to WGS from the same sample (see an example in **Supplementary Fig. 1)**.

### Copy number features associated with clinical parameters

Using a discovery cohort of 134 tumor-normal pairs, we conducted WGS analysis and identified CNAs. The cohort consisted of Stage III CRCs originating from patients that were diagnosed between 2001 and 2015 **(Table 1)**. We examined demographic (*gender, ethnicity, age*) and relevant clinical variables (*treatment, sideOfColon, cancerGrade, recurrence, BMI, smokingStatus)* associated with overall survival. The right side of colon was defined as any of “ascending colon”, “appendix”, “cecum”, “hepatic flexure”, “transverse” and the left side of colon was defined as any of “descending colon”, “rectosigmoid junction”, “sigmoid colon”, “splenic flexure”.

**Table 1.** Summary statistics and multivariate cox regression results for the Stage III colorectal cancer discovery cohort.

| Discovery Cohort (n=134) | Patient Characteristics | Count (%) / Mean (Range) | Hazard Ratio | 95% CI | p-value | significance |
| --- | --- | --- | --- | --- | --- | --- |
| **Age** | At Diagnosis | 73 (22-93) | 0.997 | (0.9715, 1.0034) | 0.795 | n.s. |
| **Gender** | Male (Reference) | 66 (49%) |  |  |  |  |
|  | Female | 68 (51%) | 0.935 | (0.5523, 1.0701) | 0.801 | n.s. |
| **Ethnicity** | White (Reference) | 128 (96%) |  |  |  |  |
|  | Hispanic | 2 (1%) | 1.236 | (0.7066, 2.1606) | 0.458 | n.s. |
| **Smoking Status** | Smoker (Reference) | 36 (27%) |  |  |  |  |
|  | Nonsmoker | 92 (69%) | 0.631 | (0.3637, 1.0951) | 0.102 | n.s. |
| **Body weight** | BMI | 29 (18-60) | 1.001 | (0.9487, 1.0552) | 0.984 | n.s. |
| **Treatment** | Chemotherapy (Reference) | 74 (55%) |  |  |  |  |
|  | Refused / Not recommended | 43 (32%) | 0.416 | (0.2393, 0.7249) | 0.002 | ** |
| **Tumor Grade** | High | 58 (43%) | 1.271 | (0.7667, 2.1082) | 0.352 | n.s. |
|  | Low (Reference) | 76 (57%) |  |  |  |  |
| **Tumor Side** | Right Colon (Reference) | 90 (72%) |  |  |  |  |
|  | Left Colon | 44 (28%) | 0.576 | (0.3423, 0.9707) | 0.038 | * |
| **Recurrence** | None | 50 (37%) | 0.652 | (0.3579, 1.1892) | 0.163 | n.s. |
|  | Recurrence (Reference) | 72 (54%) |  |  |  |  |
| **Overall Survival** | >5-year from diagnosis | 16 (12%) |  |  |  |  |

<sup>1</sup>Statistical significance is based on the fitted multivariate cox model (Eq. 1). n.s.: not significant, \*: P<0.05, \*\*: P<0.01.
<sup>2</sup>Treatment is any treatment received after surgical resection. Chemotherapy: if received any forms of 5FU, Folfox, or Capecitabine
<sup>3</sup>Missing data for each variable was <13%.

Among these Stage III CRC patients, 74 (55%) received adjuvant chemotherapy – treatment occurred with either 5-fluoruracil-based **(5-FU)** or capecitabine regimen. Patients undergoing adjuvant chemotherapy had a statistically significant improved survival as expected ^9, 10^, with a relative reduction of death risk >58% (HR= 0.416, 95% CI=[0.239, 0.724], P= 0.00195). These patients had a median overall survival of more than a year longer than that of patients who did not receive any therapy.

Stage III CRC patients with a primary tumor located in the left colon had a statistically significant improved survival with a relative reduction of death risk >42% (HR= 0.576, 95% CI= [0.342, 0.971], P= 0.0382). These patients had a median overall survival that was approximately one-half year (181 days) longer than that of patients had tumor occurred to the right of the colon.

### Analysis of recurrent arm-level CNAs

We observed extensive focal- and arm-level CNAs among the Stage III CRC discovery cohort **(Fig. 3)**. As has been widely reported and is seen with samples exhibiting CNAs, >85% of CRC are CIN phenotype and this percentage is even higher for advanced CRCs for CIN’s known to be associated with poor prognosis. The Gistic2 analysis revealed that more than half of the samples showed CNA gains of chr7, chr8q, chr13q and chr20q. And almost a half of the samples had CNA loss in chr17p and chr18.

**Figure 3.**
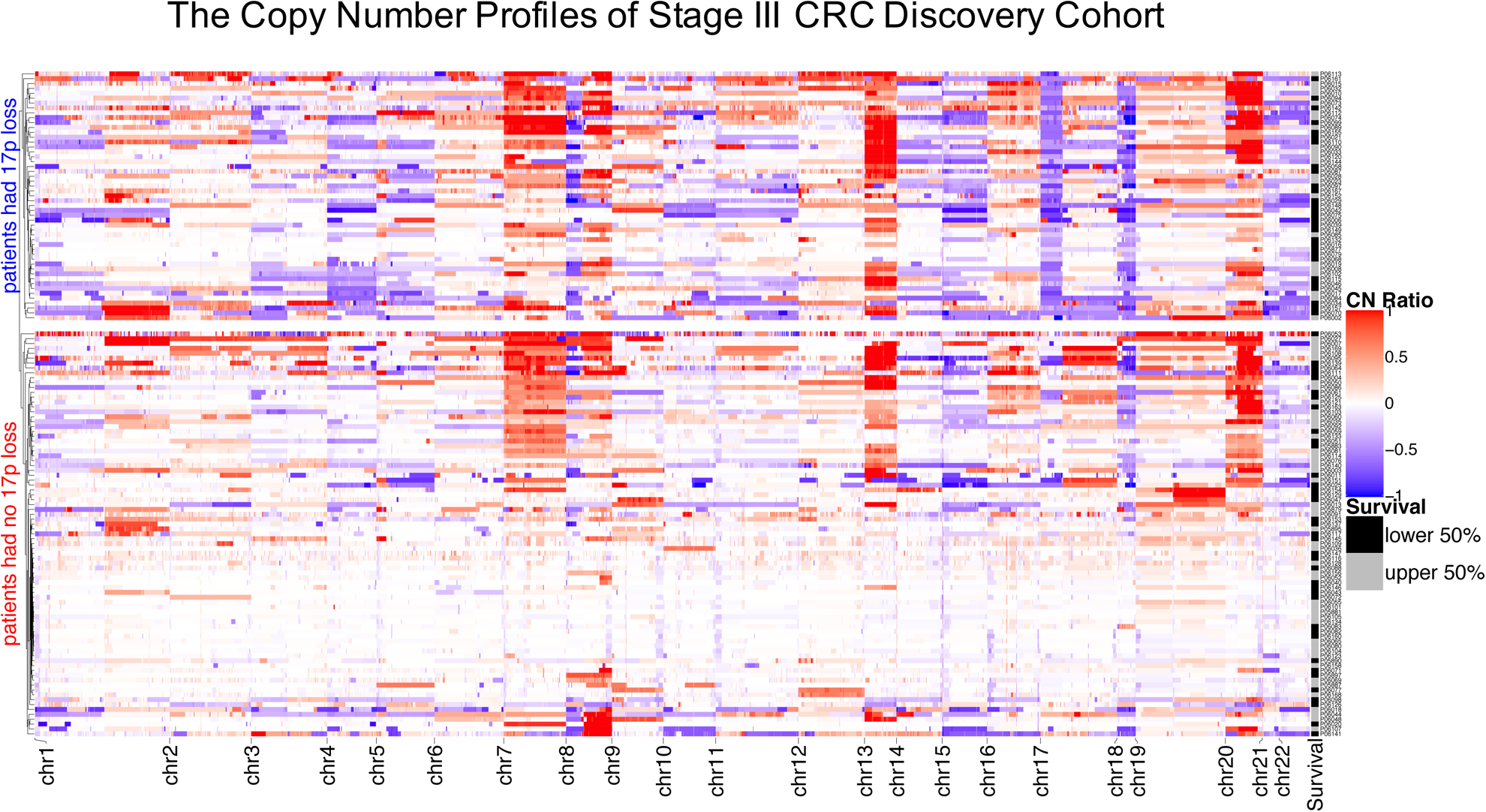
Copy number. Copy number ratios (CNR) are shown for upper split panel: patients had chr17p loss; and lower split panel patients had no chr17p loss – all based on Gistic2 calls. Row color coding: black for shorter survival patients (the lower 50%) and grey for longer survival patients (the upper 50%).

The frequency of recurrent CNA gains or losses identified in the discovery cohort were concordant with the frequency observed in the TCGA stage III validation cohort. With a q-value of <0.25 and a 99% confidence as our statistical thresholds (*Gistic2*), we identified 21 arm-level CNAs that were statistically significant **(Fig. 4)**. Seventeen CNAs were also significantly recurrent in the Stage III TCGA cohort. These included amplifications of chr1q, chr7, chr8q, chr13q and chr20, and deletions of chr1p, chr4, chr5q, chr15q, chr17p, chr18, chr21q and chr22q. Overall, the concordance of statistically significant recurrent arm-level CNAs between the two cohorts was >81%.

**Figure 4.**
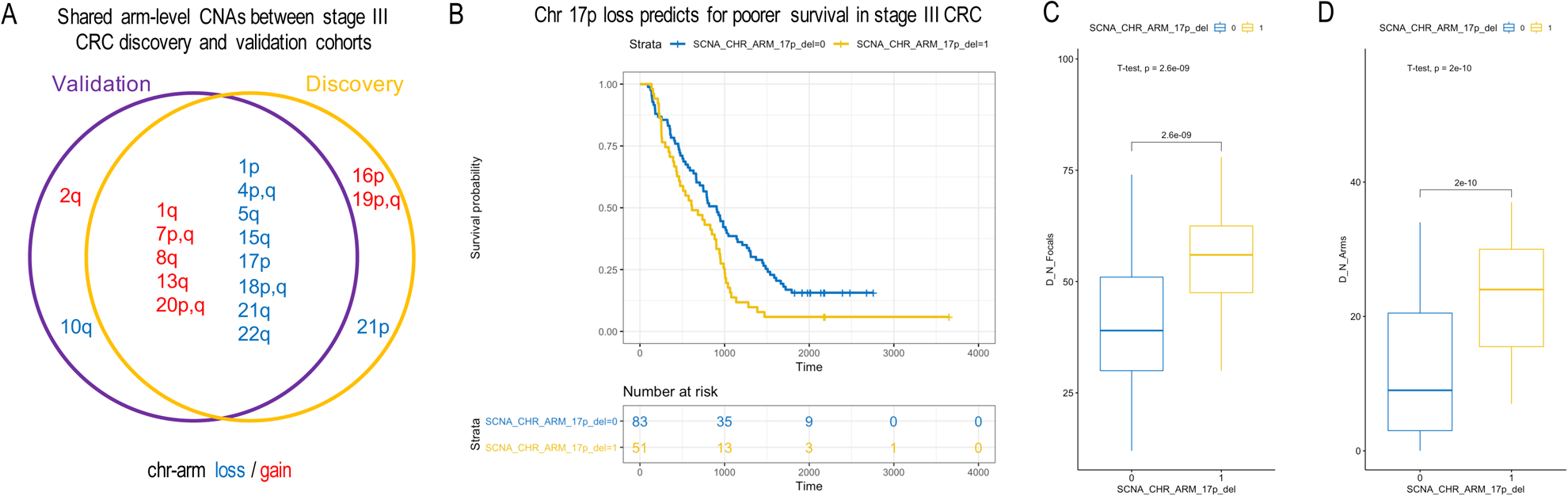
Arm-level chr17p loss predicts for poorer survival in Stage III CRC. **(A)** Venn diagram for shared arm-level CNAs between the discovery and TCGA validation cohorts. **(B)** The Kaplan–Meier plots of the Stage III CRC discovery cohort as stratified by patients’ status of carrying the chr17p arm loss (SCNA_CHR_ARM_17p_del=1 for yes, otherwise 0). Also shown are box plots for comparing **(C)** number of focal CNAs (*D_N_Focals*) and **(D)** number of arm-level CNAs (*D_N_Arms*) between patients carrying or not carrying the chr17p loss.

### Chromosome arm 17p loss is associated with poor overall survival

We applied lasso-regularized multivariate cox regression to select for significant arm-level events predictive of patients’ overall survival **(Methods)**. We identified the loss of chr17p as the single and most significant arm-level event associated with overall survival. This significance was apparent using a non-zero regression coefficient. By our final multivariate model **(Eq. 2)**, the loss of chr17p was a significant predictor of lower overall survival (P=0.0160, HR=1.706, 95% CI=[1.104-2.635]). Any patient carrying 17p deletion was associated with a 68.8% increased risk of death. An increased risk was also clear by examining the resulting Kaplan-Meijer curves when patients were stratified by their 17p loss status **(Fig. 3)**. Other factors including gender, ethnicity and age at diagnosis were not associated with overall survival. After adjusting for 17p-loss, the patients having CRC occurred to the left side of colon still had a relative reduction of death risk >33%, however, the effect ceased to be statistically significant with a marginal P=0.0727, likely a sensitivity limit imposed by the sample size. Nonetheless, patients receiving adjuvant therapy, i.e. *treatment*, (P=0.002, HR=0.478, 95% CI= [0.299-0.765]) were benefitting from a significant reduction of risk of death (52.2%) after adjusting for 17p-loss. This statistically significant difference in survival is also visible as an enrichment of shorter survival patients within the strata of carrying 17p-loss **(Fig. 3)**.

For our validation analysis, we employed the same multivariate cox model to the TCGA cohort data. Using this completely independent data set, we again confirmed that the loss of 17p was associated with poor survival (P=0.0126, HR=2.357, 95% CI=[1.202-4.621]). A patient with Stage III CRC and a 17p deletion had a 135% increased risk of death in the TCGA cohort. No other genomic, demographic and clinical factors were significant, including gender, ethnicity and age at diagnosis, except for *treatment* (P= 0.00141, HR=0.322, 95% CI=[0.161-0.646]). Receiving adjuvant chemotherapy treatment was associated with a 67.8% improvement of survival, a result that agreed with our findings from the discovery cohort and previous clinical trials.

Using our discovery cohort, we examined whether patients treated with adjuvant CRC patients (N = 69) had a lower overall survival based on 17p copy number. A 17p-loss was associated with an increased death risk at HR=1.457, CI=[0.855,2.482], P=0.166. Thus, a trend towards lower survival in the setting of adjuvant therapy was noted but the sample size was too small to reach statistical significance.

### Chromosome arm 17p loss is associated with increased chromosomal instability

*TP53*, a critical tumor suppressor involved in genome stability, is located on the 17p arm. Point mutations involving *TP53* were associated with poor survival in colorectal cancers ^11^, as indicator of TP53 loss-of-function ^12^. We compared both the focal- and arm-level CIN, as represented by the total counts of CNAs, between patients carrying or not carrying the 17p loss **(Fig. 4)**. We confound consistently higher CIN scores in patients carrying the 17p loss (one sided T-test, P=2.6e-9 for focal and P=2e-10 for arm-level CIN, respectively). This increase is notable in **Fig. 3**, where almost all patients had chr17p loss had significant copy number changes globally, indicative of a CIN subtype.

The loss of 17p thus could be an indicator for the loss of TP53 function which is known to contribute to CIN in CRC. *Vogelstein et al*. theorized that the *TP53* alteration occurs at relatively later stage of colon cancer and is responsible for promoting tumor invasion to surrounding normal tissue ^13^. Our observation of extensive presence of 17p loss in Stage III CRCs provides additional supporting evidence to this conclusion.

## DISCUSSION

This study represents one of the most comprehensive copy number analysis of Stage III CRC with the benefit of using WGS and its associated complete genome coverage and higher resolution. With these WGS results, we identified stage-specific CNA prognostic markers for Stage III CRC. Our analysis identified the loss of chromosome 17p arm, spanning *TP53*, as a potential biomarker for poor survival in Stage III CRC. Our results were independently validated by the TCGA cohort. Patients with 17p/TP53 deletion in their CRC tumors have 1.6 times relative death risk in general, as compared to those tumors which do not. Previous studies of CRC have used cohorts with mixed clinical stages. For example, nearly all of the studies included a higher number of CRCs from Stage I and II patients compared to Stage III and IV patients. Furthermore, nearly all the previous studies did not conduct an independent validation using a separate population of CRC patients with an independent validation study. The majority of these studies used low resolution molecular methods with reduced sensitivity for CNAs.

Interestingly, 17p loss has been previously reported to be prognostic marker for poor survival in other cancers, including brain tumors ^14-16^, bone tumor ^17^, periampullary cancer ^18^, pancreatic cancer ^19^, leukemia ^20^, and in CRC ^21^ evaluated with FISH. These results suggest that 17p loss may be generally useful for predicting patient outcomes. Other large-scale copy number events have been previously identified as prognostic marker for colorectal cancers, *e.g*. 18q deletion for stages II and III colon cancers ^22, 23^, which also lend support to the argument that CNA profiling may be useful for CRC management.

We identified that chromosome arm 17p CNAs occurred consistently in both the discovery and validation cohorts. Minor differences were noted between the two cohorts, as the discovery cohort had chr16p and chr19 amplifications and chr21p deletion while the validation cohort had chr2q amplification and chr14q deletion. These minor differences are likely attributable to the limited cohort size such that not enough samples were available in either cohort for determining the statistical significance of recurrent events presented at lower frequency. It may also reflect the population specific genetics for CRC progression – additional studies will be required to clarify these differences.

We also detected a trend for 17p/*TP53* loss as a predictive biomarker for poorer adjuvant chemo-therapy response in Stage III CRC. Although the finding did not achieve statistical significance due to the small cohort size, there are several pieces of additional evidences in the literature for consideration. For example, a recent publication, *Oh et al*. ^24^ has found a low-expression of TP53 protein was associated with poor cancer-specific survival in Stage III and high-rick Stage II CRC patients (N=621) who were treated with oxaliplatin-based adjuvant chemotherapy. Additionally, using the TCGA cohort, we found *TP53* copy number loss were significantly associated with lower mRNA expression level (P<1e-15, one tailed T-test on Z-normalized mRNA expression levels, see **Figure S3**). All together, these findings suggested that the loss-of-function of TP53 protein, as genetically determined by the focal *TP53* gene loss or arm-level chr17p loss, has important prognostic value for late stage CRC patients receiving adjuvant therapies.

To explain the effect of 17p loss, a likely mechanism is increased chromosomal instability, which was observed co-occurring with 17p loss. We analyzed the association between focal and arm-level chromosomal instability and 17p loss and found they were significantly associated with each other. In addition, other studies have shown that 17p loss-of-heterogeneity **(LOH)** was correlated with CRC’s metastatic potential ^25^. Similar findings were reported for other cancer types like brain tumor ^26^ and esophagus cancer ^27^. It has been reported that allelic loss of 17p allelic loss was highly correlated with *TP53* mutations ^28^. All these findings suggest that the loss of 17p is directly related to higher CIN.

There is broad interest in determining which CNAs indicate poor prognosis for individuals with Stage III CRC. We leverage the generally higher resolution of WGS analysis. WGS using a low average sequence coverage is now competitive or at an edge to microarray for both performance and economic reasons. Performance-wise, low-coverage WGS consistently showed high concordance with microarray and high-coverage WGS in our analysis. It has less noise and bias as compared to WES-based results. At 2-4x coverage, WGS provides thousands of read pairs per 100kb segment, a substantive amount enough to enable sensitive CNA detection. This provides an improved resolution compared to SNP microarrays in which there are approximately 60 probes per 100kb segment.

Identifying copy number variation with high coverage WGS data have been studied extensively in basic and clinical research settings ^29-31^. For example, in a recent systematic benchmark, *Trost et al* reported good performance of using >20x WGS data for identifying small-scale CNVs (1 – 100kb) ^32^. Our study demonstrated the feasibility and robust performance of lower coverage WGS for profiling large-scale focal and arm-level CNAs (>100kb), as an alternative to microarray and WES.

As WGS studies become less expensive, we foresee that in the future low-coverage WGS may prove to be replacing clinical microarray testing for cancers ^33^, developmental disabilities, congenital anomalies ^34-36^, autism spectrum disorder ^37^, and many other genetic diseases ^29^. Citing the benefits of WGS, a recent study compared the performance of low-coverage WGS versus microarrays on rare and undiagnosed cases. The conclusion of this study was that robust identification of CNVs was highly feasible with low-coverage WGS ^38^. In another study, low-coverage WGS also found successful application in preimplantation genetic diagnosis of monogenic disease ^39^.

## METHODS

### Discovery cohort ascertainment

The Institutional Review Boards **(IRB)** from Stanford University and Intermountain Healthcare approved the study. A total of 134 patients were recruited through the Intermountain Cancer Center (St. George, Utah, USA). Selection criteria involved those diagnosed with Stage III CRC in 2001-2015. We excluded patients who survived less than 90 days after the initial diagnosis and died from non-cancer causes. We collected relevant clinical information from patient medical records, including age of diagnosis, gender, ethnicity, body mass index **(BMI)**, and smoking status **(Table 1** and **Supplementary Table 1)**.

### DNA extraction from clinical samples

We collected matched primary colorectal adenocarcinoma tumor and normal colon tissue samples from each patient **(Fig. 1)**. All samples were determined to have greater than 60% tumor content in pathology review. We used a two-millimeter punch from a tumor or normal FFPE tissue block. The DNA was isolated from tissue using the Maxwell-16 and Promega-AS1030 DNA purification kit (Promega, Wisconsin, USA). The genomic DNA was quantified via the Qubit (Thermo-Fisher Scientific, Massachusetts, USA) and quality assessment was performed with the LabChip GX (PerkinElmer, Massachusetts, USA).

### Sequencing

For sequencing library preparation, 500 nanograms of DNA from each sample was sheared using a Covaris E220 (Covaris, Massachusetts, USA) with microtube plates and following parameters: intensity level of five, duty cycle of 10%, cycles per burst of 200, and treatment time of 55 seconds. The DNA was then purified with a 0.8X AMPure XP (Beckman-Coulter, California, USA) bead cleanup to maintain a large insert size for sequencing. We used this total yield of purified DNA for the Kapa Hyper Prep Kit for Illumina (Roche, Basel, Switzerland). The standard KAPA protocol was followed with eight cycles of PCR amplification and a 0.8X post-amplification cleanup. We used 10 base pair dual-index sequencing adapters to allow for index swapping detection.

We measured the library quality with the LabChip GX and quantity with the Qubit **(Supplementary Table 2)**. The libraries were pooled and sequenced on an Illumina MiSeq (Illumina, California, USA) for paired-end 300 basepair reads. The sequencing libraries were repooled and normalized based on the MiSeq data before paired-end 300 basepair sequencing on an Illumina NovaSeq 6000 system achieving 2-4x coverage per sample. Sequence reads were aligned to the human reference genome GRCh37/hg19 with the Burrows-Wheeler Aligner ^40^.

### Copy number segmentation

For determining which copy number segmentation tool provided accurate results on WGS from FFPE-extracted DNA, we evaluated the copy number callers, CNVkit ^41^ and BicSeq ^42^ – both are readily available as open source scripts. Segmentation involves defining the intervals that are affected by a copy number change. As test data set, we downloaded the low-coverage (∼5x) WGS and the SNP-array data of 10 randomly selected **(Supplementary Table 3)** CRC tumor-normal pairs from TCGA. Using the WGS data, we inferred the genome segments with CNVkit and BicSeq, and estimated log copy number ratio per segment **(CNR)**. For each sample, we correlated the WGS-estimated segmental CNRs to the micro-array CNRs using 100-kb genome-wide tiles. We computed the Pearson’s correlation coefficient **(PCC)** between the two set of estimates and summarized PCC over all samples by mean and standard deviation. We compared the metric difference between groups using the two-sided T-test. We also conducted a gene-based PCC analysis using the same data.

Next, we evaluated how WGS coverage reduction affects genome segmentation. We applied ∼30X high-coverage WGS analysis to eight patients with the identical protocol to low-coverage WGS. The high-coverage sequence data were down-sampled to low-coverage (∼3x) data. We performed genome segmentation using CNVkit on both the high- and low-coverage WGS data. We computed and compared the PCC metrics for high- and low-coverage CNRs based on tumor purity.

WES and targeted sequencing are other common choices for CNA analysis. We also bench-marked low-coverage WGS to high-coverage WES (∼300x) in a random subset of 10 patients. We performed genome segmentation using CNVkit on both the high-coverage WES and low-coverage WGS data. We computed and compared the intra-tile deviation metrics of estimated segmental log CNRs based on 100kb genome tiles.

### Integrated copy number analysis pipeline

We integrated CNVkit ^41^, Gistic2 ^43^, *coxph, survival* and *glmnet* ^44, 45^ packages of R into our final copy number analysis bioinformatics pipeline **(Fig. 1)**. We used the data from the genome segments inferred by CNVkit to Gistic2, a cohort CNA caller. We ran Gistic2 with the following arguments: *“-refgene hg38.UCSC.add_miR.160920.refgene.mat -maxspace 10000 -ta 0.1 -td 0.1 -qvt 0.25 -broad 1 -brlen 0.7 -twoside 1 -conf 0.99 -genegistic 1 -armpeel 1 - savegene 1 -res 0.05 -smallmem 1 -js 4”*. We set the noise cut-off for both deletion and amplification to 0.1. Coupling CNVkit with Gistic2 ^43^ enabled us to identify recurrent arm and focal-level CNAs with statistical significance. We also integrated and *ggplot2, survminer, ggpubr, inferCNV* R packages ^46^ for data visualization.

To control for false positives, we identified error-prone CNA regions that demonstrated a high level of CNV background noise using normal DNA samples which had no somatic copy number changes. We ran genome segmentation and CNA calling on all normal DNA samples 10 times and each time with one random normal sample as reference. We compiled all of the CNA calls and identified regions that demonstrated copy number changes in >10% of samples in each run for >5 runs. These changes were likely the result of false copy number calls that were specific to FFPE-extracted DNA, amplification bias or sequencing artifacts. We filtered out the false positive segments from those noisy regions before conducting the Gistic2 analysis.

### Stage III-specific biomarker discovery

We first fitted a multivariate cox model with relevant clinical and demographical covariates to identify any such variables was associated with survival. The full model is as follows:

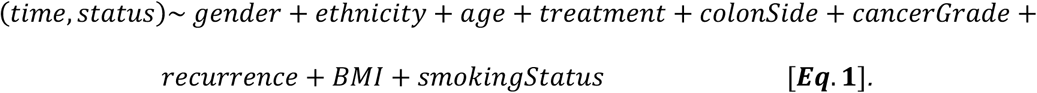

Next, we created binarized CNA variables for each arm-level deletion or amplification (e.g. chr1p_del_ or chr1p_amp_ for chr1) using the *Gistic2* output. The variable is coded one if the CNA amplitude exceeds the noise cut-off, otherwise zero. We fitted a multivariate cox regression model with lasso-regularization to select for candidate CNA biomarkers ^47^, including all the 88 arm-level CNA variables, gender, ethnicity, age and treatment.

Finally, we tested the lasso-selected candidate CNA variables’ significance by the following multivariate cox regression model:

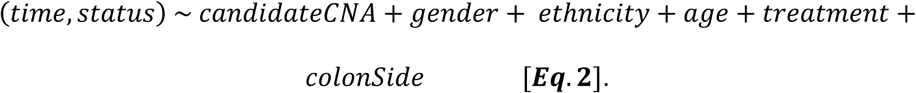

The resulting p-values were adjusted by the Bonferroni correction and significant results were declared with Bonferroni-adjusted P<0.05.

### Biomarker validation

CNA biomarkers were validated with an independent cohort of 187 Stage III CRCs from TCGA **(Supplementary Table 4)**. We downloaded the overall survival time, survival status, SNP-array based segments, and other clinical data for these patients in the validation cohort. The genome segmentation data was formatted for Gistic2 analysis. With this independent validation dataset, we tested the same multivariate model **(Eq. 2)**, including these candidate CNA markers in question. A candidate marker was declared statistically significant only if the Bonferroni-adjusted P<0.05.

### Chromosomal-instability analysis

We measured CIN at both focal and arm-level by counting the total number of such events sample-wise. We denoted the arm-level CIN by *D_N_Arms*, which is the total number of arm-level CNAs per sample. Similarly, we denoted focal-level CIN by *D_N_Focals*, which is the total number of focal-level CNAs per sample. We applied a T-test to determine if any significant difference in CIN in patient groups as stratified by chr17p_del_ status using the *D_N_Focals* and *D_N_Arms* measures.

## AUTHOR CONTRIBUTIONS

LDN, PVH and HPJ designed the study. TB, DSH and GF collected the clinical information. MK and CWB conducted the sequencing. LCX, HL, SMG, JMB and SUG performed bioinformatics analysis. LCX performed the statistical analysis. LCX, MK, PVH, HPJ and LDN wrote the manuscript. All authors edited, revised, read and approved the manuscript for submission.

## ADDITIONAL INFORMATION

### Acknowledgements

This work was supported by the following grants from the NIH: P01HG000205 to MK, JMB and HPJ; 1U01CA15192001-A1 to HJL and HPJ; 1U01CA176299 to HJL and HPJ, HG006137-07 to LCX and HPJ. An award from Intermountain Healthcare supported LCX, HPJ and MK. HPJ was also supported by a Research Scholar Grant (RSG-13-297-01-TBG) from the American Cancer Society. HPJ also received grant and research support from the Clayville Foundation. LCX was also supported by a research grant from the Innovation in Cancer Informatics Fund and a postdoc fellowship grant from American Cancer Society (PF-18-184-01-TBG). LDN was supported by NIH/NCI (5K08CA166512), and the Conquer Cancer Foundation (Young Investigator Award), the Gastric Cancer Foundation, and the Carl Kawaja Foundation.

### Data availability

The copy number segmentation and survival data of the 134 stage-III colorectal cancer cohort are available with the supplementary files associated with this paper. The copy number segmentation and survival data of TCGA colorectal cancer cohort are available from the National Cancer Institute’s Genomic Data Commons (https://gdc.cancer.gov).

### Ethics approval and consent to participate

The Institutional Review Boards from Stanford University and Intermountain Healthcare approved the study. The study was performed in accordance with the Declaration of Helsinki.

### Consent for publication

Not applicable.

## SUPPLEMENTARY FIGURES

**Supplementary Figure 1.**
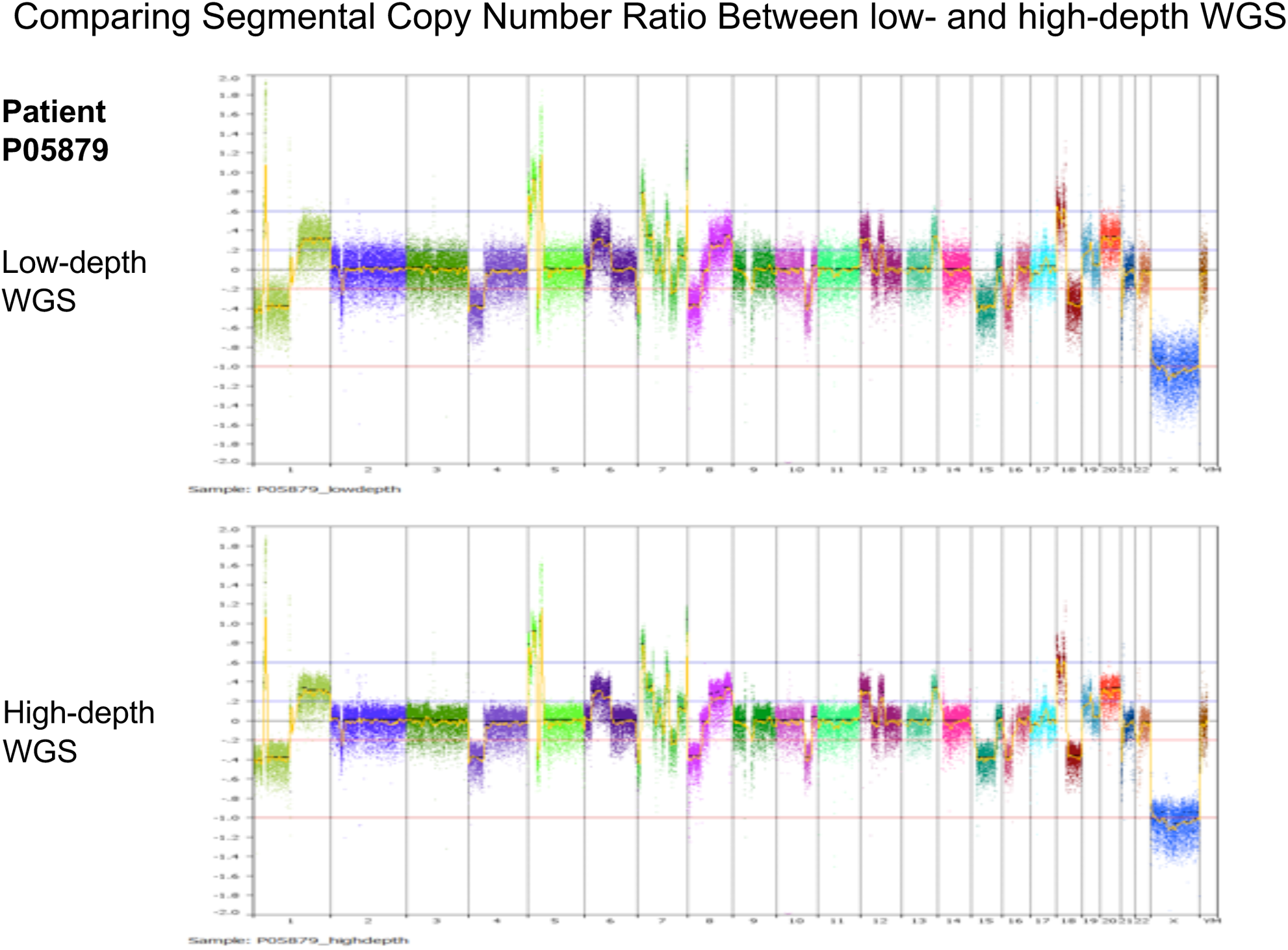
A shoulder-to-shoulder view of the segmentation tracks estimated from the low-coverage WGS and WES platforms from the same sample. Y-axis was drawn in the same scale. This suggests WES-derived segments are significantly more variable as compared to low-coverage WGS.

**Supplementary Figure 2.**
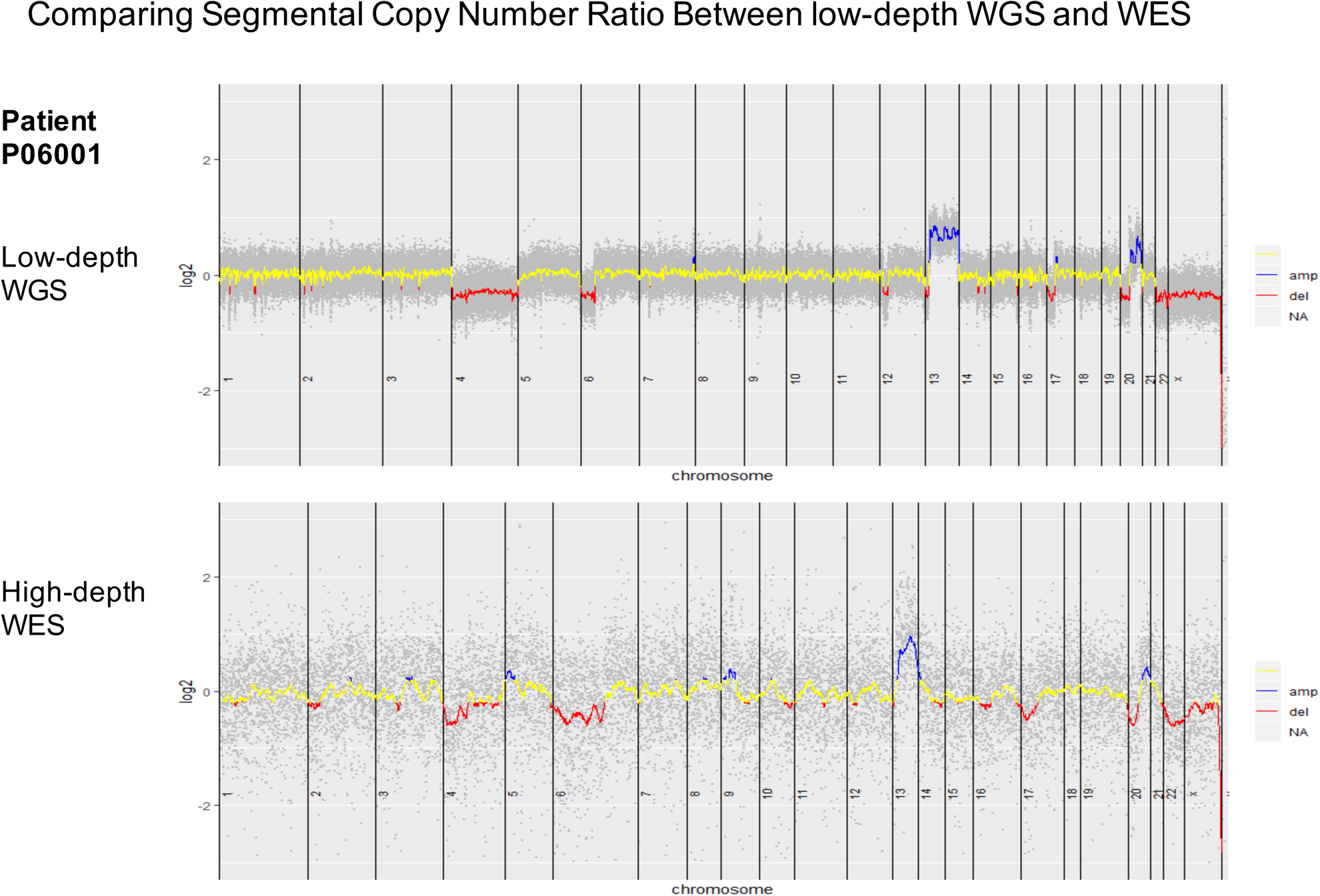
A shoulder-to-shoulder view of the segmentation tracks estimated from the low-coverage and high-coverage WGS platforms from the same sample. Y-axis was drawn in the same scale. This suggests for this high purity tumor sample, the low-coverage WGS-derived segments are highly accordant to high-coverage WGS’s.

**Supplementary Figure 3.**
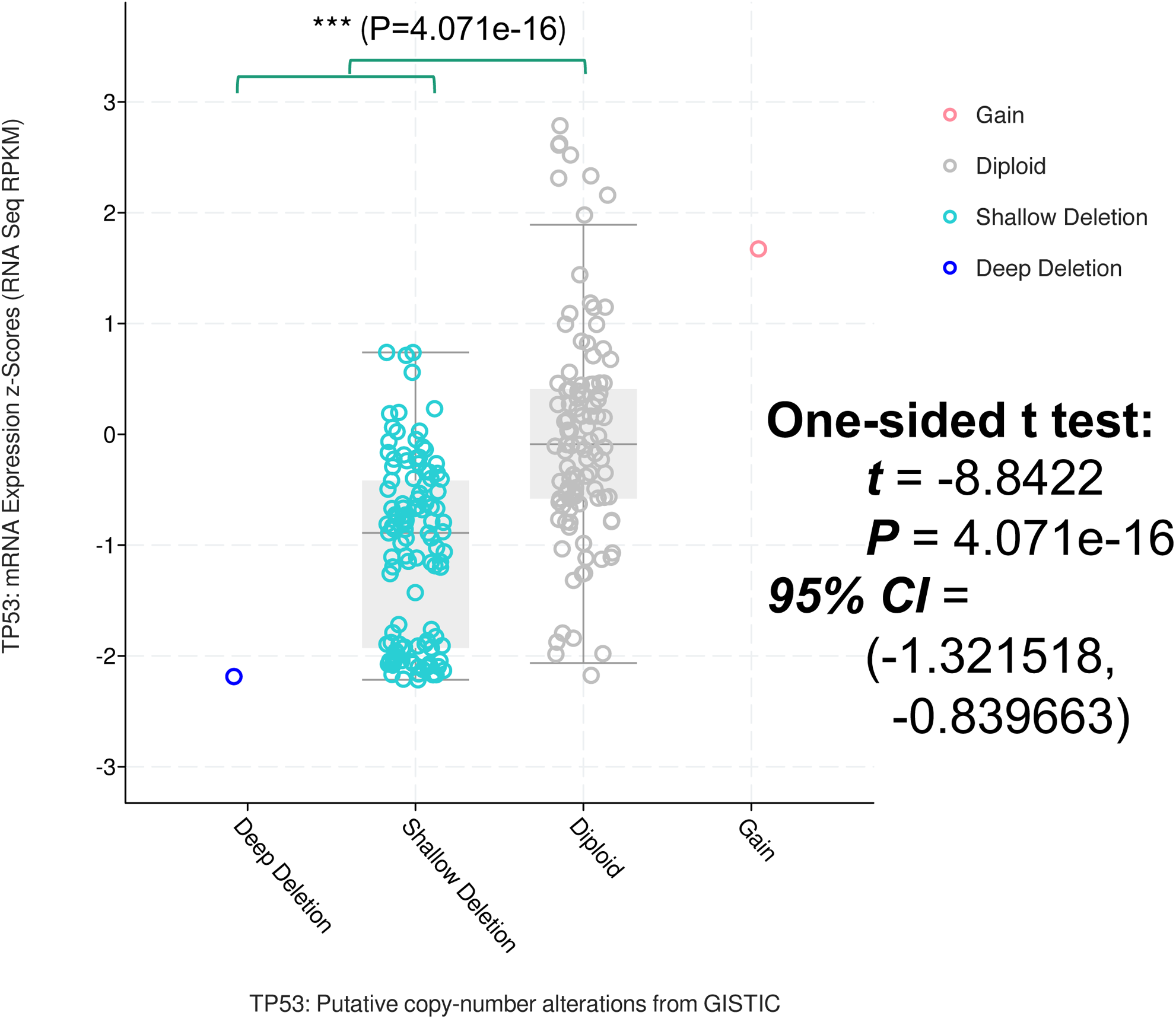
mRNA expression difference between patients with and without TP53 deletion in the TCGA cohort. This suggests TP53 deletion is associated with significantly reduced mRNA expression level of TP53.

